# Why did some practices not implement new antibiotic prescribing guidelines on urinary tract infection? A cohort study and survey in NHS England primary care

**DOI:** 10.1101/355289

**Authors:** Richard Croker, Alex J Walker, Ben Goldacre

**Affiliations:** Honorary Research Fellow, EBM DataLab, Centre for Evidence Based Medicine, Nuffield Department of Primary Care Health Sciences, University of Oxford, OX2 6GG; Researcher, EBM DataLab, Centre for Evidence Based Medicine, Nuffield Department of Primary Care Health Sciences, University of Oxford, OX2 6GG

**Author notes:** Dr Ben Goldacre (corresponding), Director, EBM DataLab, Senior Clinical Research Fellow, Centre for Evidence Based Medicine, Nuffield Department of Primary Care Health Sciences, University of Oxford, Radcliffe Observatory Quarter, Woodstock Road, Oxford OX2 6GG.

**Keywords:** Prescribing, Urinary tract infection, antibiotics, antibiotic resistance

## Abstract

**Objectives:** To describe prescribing trends and geographic variation for trimethoprim and nitrofurantoin; to describe variation in implementing guideline change; and to compare actions taken to reduce trimethoprim use in high- and low-using Clinical Commissioning Groups (CCGs).

**Design:** A retrospective cohort study and interrupted time series analysis in English NHS primary care prescribing data; complemented by information obtained through Freedom of Information Act requests to CCGs. The main outcome measures were: variation in practice and CCG prescribing ratios geographically and over time, including an interrupted time-series; and responses to Freedom of Information requests.

**Results:** The amount of trimethoprim prescribed, as a proportion of nitrofurantoin and trimethoprim combined, remained stable and high until 2014, then fell gradually to below 50% in 2017; this reduction was more rapid following the introduction of the Quality Premium. There was substantial variation in the speed of change between CCGs. As of April 2017, for the 10 worst CCGs (with the highest trimethoprim ratios): 9 still had trimethoprim as first line treatment for uncomplicated UTI (one CCG had no formulary); none had active work plans to facilitate change in prescribing behaviour away from trimethoprim; and none had implemented an incentive scheme for change in prescribing behaviour. For the 10 best CCGs: 2 still had trimethoprim as first line treatment (all CCGs had a formulary); 5 (out of 7 who answered this question) had active work plans to facilitate change in prescribing behaviour away from trimethoprim; and 5 (out of 10 responding) had implemented an incentive scheme for change in prescribing behaviour. 9 of the best 10 CCGs reported at least one of: formulary change, work plan, or incentive scheme. None of the worst 10 CCGs did so.

**Conclusions:** Many CCGs failed to implement an important change in antibiotic prescribing guidance; and report strong evidence suggesting that CCGs with minimal prescribing change did little to implement the new guidance. We strongly recommend a national programme of training and accreditation for medicines optimisation pharmacists; and remedial action for CCGs that fail to implement guidance; with all materials and data shared publicly for both such activities.

## Introduction

### Background

Urinary-tract infections (UTIs) are a common presentation in primary care, accounting for between 1-3% of all GP consultations.^1^ Incidence of UTIs increases with age: a recent cohort study found that 21% of patients over 65 had at least one infection in a 10 year period.^2^ Until recently, trimethoprim was the most commonly prescribed antibiotic used for empirical treatment of uncomplicated UTI, with approximately 3.9 million prescriptions dispensed in 2013.^3^ However there is now growing concern about trimethoprim resistance. In 2016 approximately 34% of samples tested in laboratories were trimethoprim resistant, compared with 3% for nitrofurantoin resistance.^4^ This increase in resistance is thought to have contributed to increased incidence of *Escherichia coli* bacteremia.^4^ Reducing the harm from inappropriate management of UTI is a priority for the NHS.^5^

In 2014 Public Health England (PHE) revised their guidance on treatment for uncomplicated UTI in primary care, recommending nitrofurantoin as first line treatment, and trimethoprim only where there is low risk of resistance (defined as “younger women with acute UTI and no risk”).^6^. In September 2016 the “Quality Premium” (QP) for Clinical Commissioning Groups (CCGs)^5^ set out financial incentives for reducing the proportion of trimethoprim prescribed, and reducing gram-negative bloodstream infections. Achievement was worth approximately £0.85 per CCG population-area patient, disbursed directly to the CCG if they achieved a 10% reduction over 12 months compared to the previous year. The same amount was available for the reduction of 10% of trimethoprim prescriptions for patients aged 70 or over. This money is given to the CCG: there is no requirement in the QP for CCGs to pass these financial incentives directly on to GP practices. These targets were for 2017-18, with a review planned for 2018-19.

Although NHS England commission NHS GP services, responsibility for the cost and quality of prescribing rests with CCGs. This is managed by Medicines Optimisation teams, hosted by CCGs or Commissioning Support Units (CSUs). Individual structures and functions vary: generally a team will include pharmacists who work with GP practices, identifying improvements in cost or clinical effectiveness, alongside nurses, technicians, and/or managers. Many CCGs also work with other stakeholders in the local health economy to produce local formularies, to promote evidence-based prescribing. Many also have “incentive schemes”, where GP practices receive financial incentives in order to facilitate improvements in specific areas of prescribing behaviour, such as those described in the QP.

In this paper we set out to: describe long-term prescribing trends for trimethoprim and nitrofurantoin; describe variation between practices and CCGs in their implementation of the new guidelines over time; to describe and map current variation at CCG and practice level; to identify the highest and lowest users of trimethoprim nationally; and to compare the actions taken by these CCGs to reduce trimethoprim use.

## Methods

### Study design

We analysed prescribing practice by conducting a retrospective cohort study in prescribing data from all English NHS GP practices and CCGs; and assessed actions undertaken by CCGs to implement new guidance by requesting information on key change activities through Freedom of Information (FOI) requests to CCGs.

### Setting and data

We extracted data from our OpenPrescribing.net database. This imports openly accessible prescribing data from the large monthly files published by the NHS Business Services Authority^7^ which contain data on cost and volume prescribed for each drug, dose and preparation, for each month, for every individual general practice and CCG in England.

### Variation between CCGs and practices in UTI treatment choices

We extracted data on all prescriptions dispensed between January 2011 and December 2017 for prescribing of trimethoprim and nitrofurantoin of any form, using the British National Formulary (BNF) codes for trimethoprim (0501080W0) and nitrofurantoin (0501130R0). Although other treatments such as cephalosporins^2^ have been used in the past for UTIs, these were excluded from our analysis as they are not covered by the guideline change nor the incentive scheme; and because they are commonly used for other indications; there is no national prescribing dataset that includes data on indication. We calculated CCG- and practice-level deciles at each month for the proportion of trimethoprim and nitrofurantoin that was prescribed as trimethoprim. These were plotted on a time series chart.

### Measuring the impact of guidance changes and financial incentives

We used interrupted time series analysis to measure the degree to which the proportion of trimethoprim prescribing changed over time after the change in PHE guidance and the start of the QP monitoring period. The analysis was used to determine the change in gradient of a regression lines before and after both interventions, as well as 95% confidence intervals for the gradient changes.

### Geographical Variation across England’s practices and CCGs

We created choropleth maps of trimethoprim prescribing proportion for three key periods of interest: the six months prior to the PHE guidance; the six months after it was released; and the first six months of the QP measurement period.

### Describing CCG-level strategies for change of prescribing behaviour

To collect data on the actions undertaken by CCGs to change prescribing from trimethoprim to nitrofurantoin we submitted a FOI request to every CCG in England, asking the information described in Appendix 1. In brief, we asked CCGs about the current status of trimethoprim and nitrofurantoin in the local formulary (if they had one); if and when this changed; whether they had any action plans for change since 2014; and whether GP practices were financially incentivised to work on these plans.

Using the OpenPrescribing.net measure “Antibiotic stewardship: prescribing of trimethoprim vs nitrofurantoin by all CCGs”^8^ we identified the 10 CCGs with the highest and lowest current proportion of prescribing of trimethoprim for the six months June-November 2017. For these CCGs we extracted structured data from the FOI responses describing whether each CCG had described change activity in each domain. We also calculated summary statistics on the prescribing behaviour for the six months prior to the PHE guidance, the six months after it was released, and the first six months of the QP measurement period.

### Software and reproducibility

Data management and analyses were performed using Python, from a data warehouse in Google BigQuery. All data, underlying code and all our tools are shared for review, and free re-use, under the MIT open license.^9^

## Results

### Variation between CCGs and practices in UTI treatment choices

Figure 1 shows trimethoprim prescribing, as a proportion of trimethoprim and nitrofurantoin prescribing, for each decile of CCG and practice over the period 2011-2017. Both practice and CCG prescribing of trimethoprim remained stable between 2012 and late 2014, when PHE guidance was amended to make nitrofurantoin first line. At this point, the proportion of trimethoprim started to slowly reduce across all deciles. From early 2017, when the QP measurement was introduced, there was a more rapid decline in trimethoprim prescribing across all deciles and extreme percentiles. Although population prescribing overall has changed to be more in line with current guidance, the level of variation between CCGs has increased since the guidance changed in 2014 (seen in the widening gap between first and ninth decile) and not yet declined to reflect a new stable consensus.

**Figure 1:**
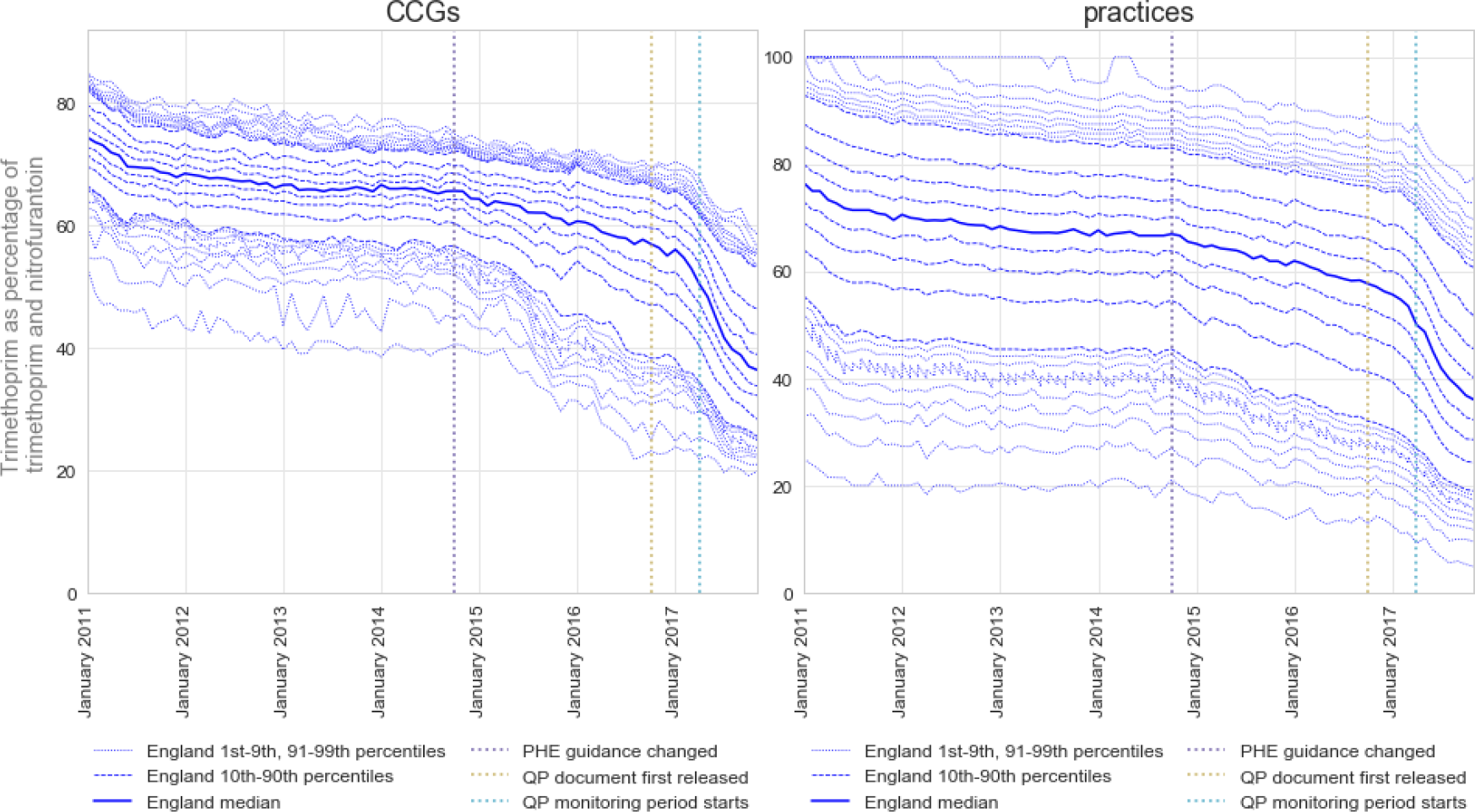
CCGs and practice decile and extreme percentile (1-9, 90-99) time trends, 2011- 2017

### Measuring the impact of guidance changes and financial incentives

Using interrupted time series analysis to assess the impact of the interventions, we determined that both the change of PHE guidance and the start of the QP monitoring period were associated with a change in gradient of prescribing proportion. However the QP monitoring period was associated with a much greater change in gradient (−1.36% per month, 95% confidence interval −1.62 to −1.10) than the PHE guidance (−0.36% per month, 95% confidence interval −0.39 to −0.33) (Figure 2).

**Figure 2:**
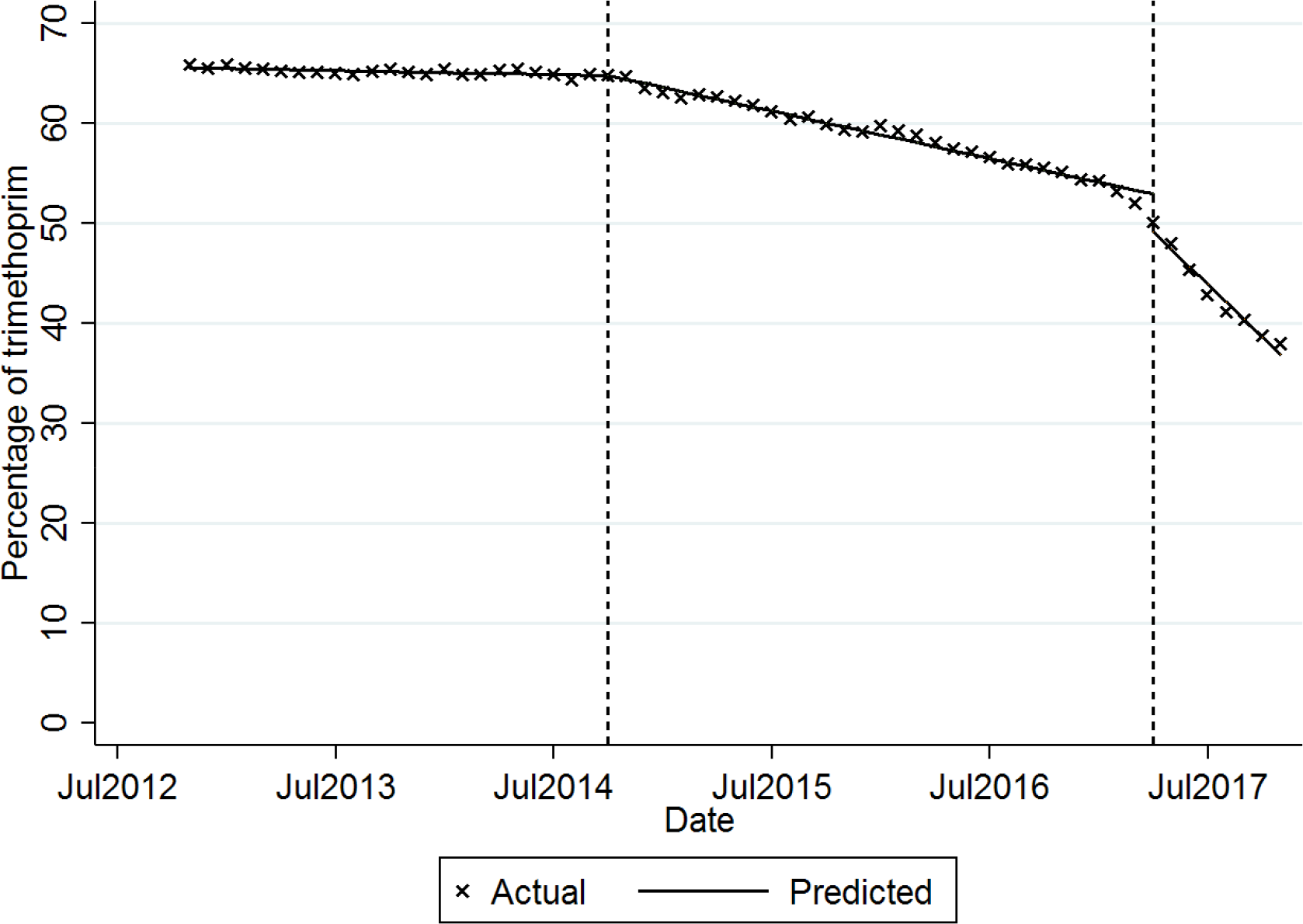
Interrupted time series analysis to determine the impact of the release of PHE guidance (left vertical line) and the QP monitoring period starting (right vertical line)

### Geographical Variation across England’s practices and CCGs

Geographical variation before, between, and after the PHE and NHS England changes is shown in Figure 3; here again it can be seen that there was little change before the introduction of the QP. Although, overall, CCGs have now reduced prescribing of trimethoprim, substantial variation remains.

**Figure 3:**
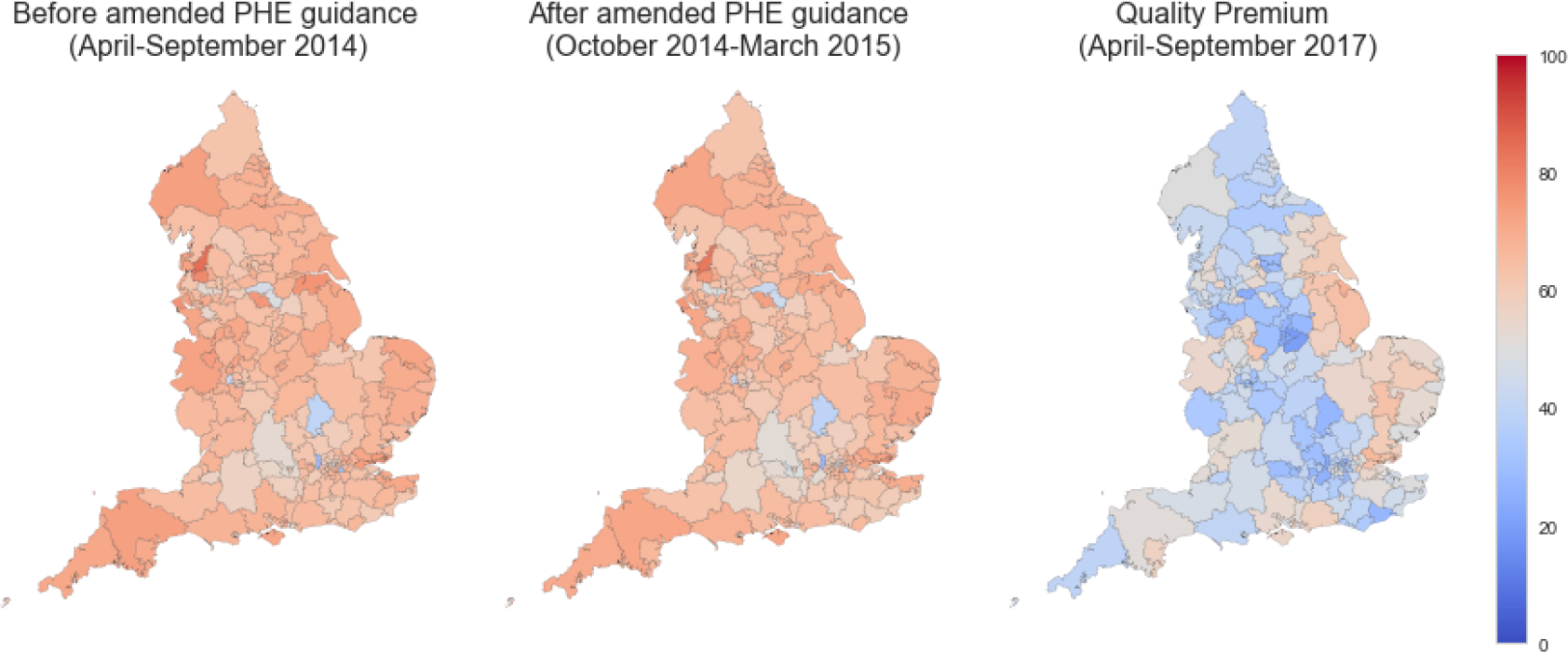
geographical variation for proportion of trimethoprim and nitrofurantoin prescribing as trimethoprim before and after PHE guidance amended, and six months after introduction of the QP

### Describing CCG-level strategies for change of prescribing behaviour

The 10 highest and lowest CCGs for trimethoprim use all replied to our FOI request. Results are summarised in Table 1 and show a pronounced difference in activity between those performing best and worst at implementing new antibiotic prescribing guidance. As of April 2017, for the CCGs with the 10 highest trimethoprim ratios: 9 still had trimethoprim as first line treatment for uncomplicated UTI (one CCG had no formulary); none had active work plans to facilitate change in prescribing behaviour away from trimethoprim; and none had implemented an incentive scheme for change in prescribing behaviour. For the CCGs with the 10 lowest trimethoprim ratios: 2 still had trimethoprim as first line treatment (all CCGs had a formulary); 5 (out of 7 who answered this question) had active work plans to facilitate change in prescribing behaviour away from trimethoprim; and 5 had implemented a financial incentive scheme for practices to facilitate change in prescribing behaviour. The change in prescribing behaviour over time for the highest and lowest current prescribers of trimethoprim at CCG level is given in Figure 4.

**Table 1:** FOI responses for 10 CCGs with highest and lowest proportion of trimethoprim to nitrofurantoin prescribing.

|  | Highest trimethoprim users (n=10) | Lowest trimethoprim users (n=10) |
| --- | --- | --- |
| *Percentage trimethoprim at baseline (April-September 2014) (range)* | 70.88%<br>(67.03%-76.55%) | 59.22%<br>(28.57%-69.39%) |
| *Percentage trimethoprim at first six months after PHE guidance amended (range)* | 69.67%<br>(66.19%-72.70%) | 56.87%<br>(27.46% - 66.58%) |
| *Percentage trimethoprim during first 6 months QP monitoring (range)* | 63.05%<br>(58.91% - 68.73%) | 24.65%<br>(19.46%-27.44%) |
| *Number of CCGs with local antibiotic formulary* | 9 | 10 |
| *Number of CCGs with trimethoprim first line treatment at April 2016* | 9 | 2 |
| *CCGs with trimethoprim first line treatment at April 2017* | 9 | 2 |
| *CCG with work plans in place at April 2016* | 0 | 5 (of 7 CCGs who answered this question) |
| *CCGs with work plans in place at April 2017* | 0 | 5 (of 7 CCGs who answered this question) |
| *CCGs with practice-level incentive schemes for 2017-18* | 0 | 5 |

**Figure 4:**
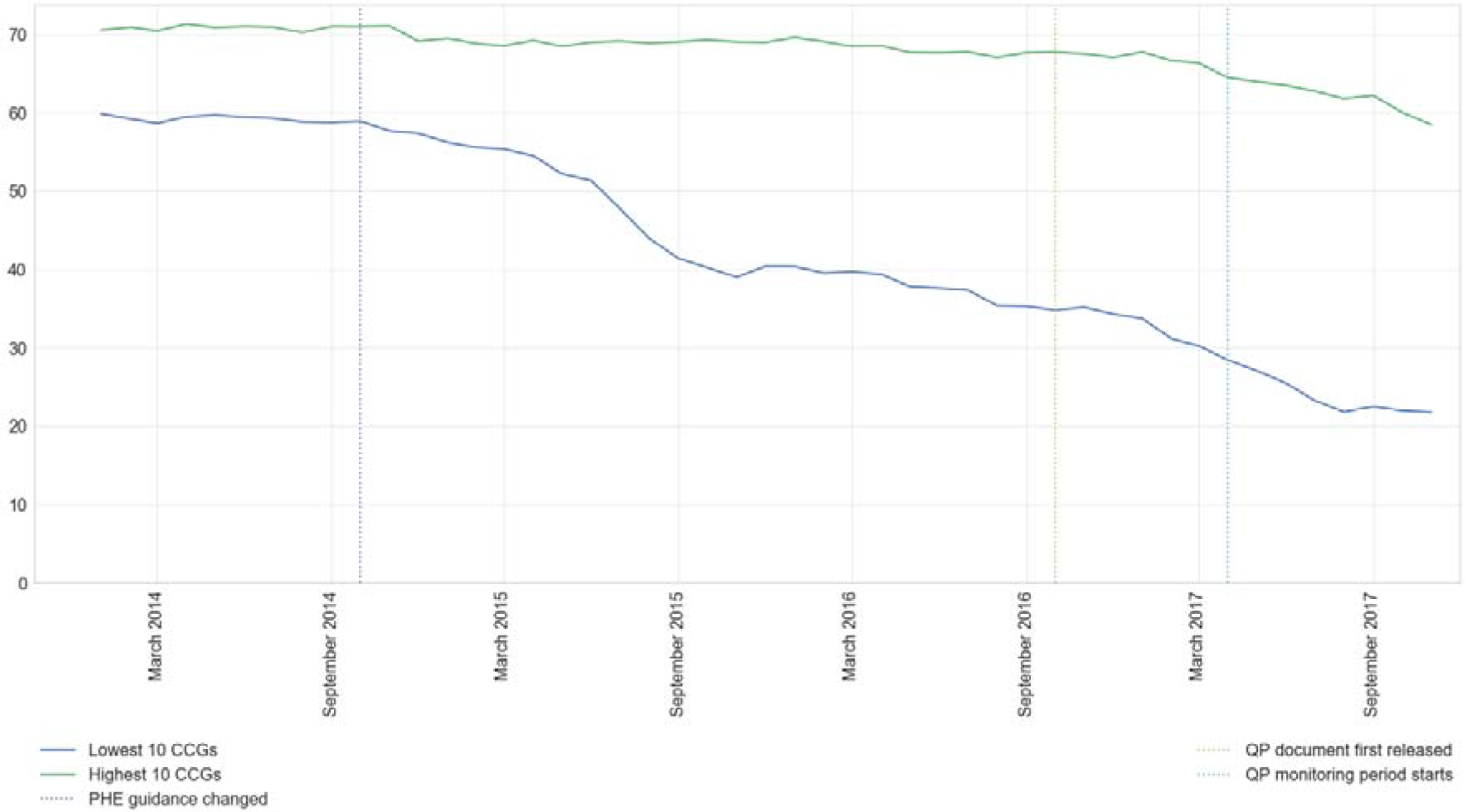
Mean proportion of trimethoprim and nitrofurantoin prescribing as trimethoprim in 10 highest and lowest CCGs, 2014-2017.

### Change over time, for best and worst prescribers

To further explore change over time we present examples of individual CCGs’ prescribing behaviour during the period 2012-2018, in the context of their practice change activity. Prescribing data for these four example CCGs is presented in Figure 5. For clarity, these four brief vignettes are intended as illustrations of change activity in organisations that implemented change successfully, and the starting point for future research, rather than definitive endorsements of individual activities.

**Figure 5 a-d:**
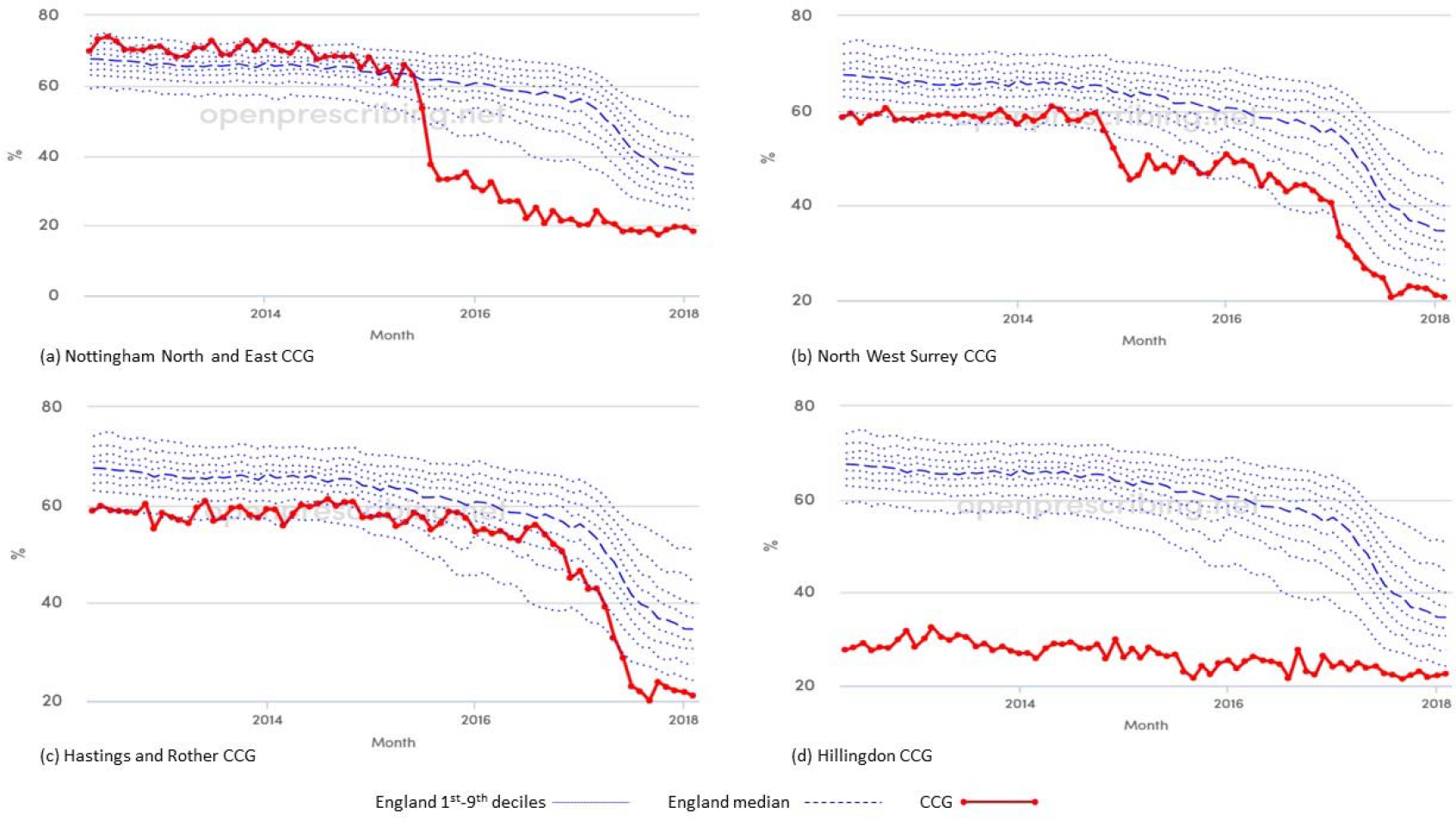
Proportion of trimethoprim prescribing 2012-2018 in: (a) Nottingham North and East CCG; (b) North West Surrey CCG; (c) Hastings and Rother CCG; (d) Hillingdon CCG.

Some CCGs, such as Nottingham North and East CCG (and other Nottinghamshire CCGs) had a relatively sudden drop in trimethoprim prescribing shortly after the PHE guidance was amended (Figure 6a). This CCG changed its antimicrobial guidelines and added a very clear message: *“Due to increasing resistance trimethoprim is NO longer recommended as empiric therapy; Nitrofurantoin is now first line treatment, however before prescribing, risk factors for resistance should be considered”.* They also specifically discussed the issue at GP practice meetings in 2015 and 2016. There was no financial incentive offered to change behaviour.

North West Surrey CCG had a different profile of change (Figure 6b), with an early reduction in 2016, but a larger reduction around the introduction of the QP. This CCG had a robust and focused action plan for its practices, including: a consultant microbiologist-led education session for GPs; funding for an audit of compliance against PHE guidance for UTI treatment; and funding for achievement of a target of 37% or below for trimethoprim as a proportion of trimethoprim and nitrofurantoin. Significant reductions in the prescribing of trimethoprim occurred after these interventions.

Hastings and Rother CCG only reduced trimethoprim use in late 2016, despite their formulary changing to nitrofurantoin as first-line treatment in March 2015 (Figure 8). They also had a simple incentive scheme for GP practices which started in April 2017, paying practices to reduce or maintain prescribing of trimethoprim and nitrofurantoin to a ratio of 1:1. The largest reductions in prescribing of trimethoprim occurred shortly after this scheme was initiated.

Hillingdon CCG has always had relatively low trimethoprim prescribing (Figure 9). Their local prescribing guidelines did not include trimethoprim, even prior to 2014. They continue to audit prescribing at practice level as part of their prescribing incentive schemes.

## Discussion

### Summary

Clinical practice around antibiotic prescribing has changed in response to guideline changes and financial incentives, but slowly and incompletely. CCGs exhibiting substantial and rapid change have implemented various change strategies. Strikingly, for CCGs exhibiting little change in antibiotic prescribing, we found no evidence for any change strategy. Only half of the 10 CCGs with the lowest trimethoprim prescribing rates have offered direct financial incentives to their prescribers, and therefore this does not appear to be necessary in order to facilitate change. However at a population level the introduction of financial incentive schemes for CCGs led to a measurably larger rate of change, compared with the change seen at the introduction of updated clinical guidance from PHE. The renewed Quality Premium for 2018-19 no longer includes the trimethoprim/nitrofurantoin target, presumably as a result of the level of change shown in 2017-18.^10^

### Strengths and weaknesses

We were able to use prescribing data to accurately show how doctors, facilitated by CCGs, responded to a change in national guidance, and to the introduction of financial incentives driven by an objective prescribing measure. Our data is complete, and covers the entire population of all NHS England GPs, rather than a sample. It is likely to be a highly accurate record of what was dispensed, as it is the basis for NHS reimbursement to pharmacies. As such, it represents the total dispensed, rather than prescribed; however there is no reason to believe patients are differentially less likely to present prescriptions for trimethoprim or nitrofurantoin to pharmacists, therefore this is unlikely to be a source of bias. As stated earlier, we excluded data on other antibiotics due to the range of conditions that they were likely to be prescribed for, and because they are not covered by any of the national prescribing policy changes; given that trimethoprim was the antibiotic previously recommended as a first line treatment, this is unlikely to have a significant impact on our findings.

A high proportion of CCGs responded to our requests under the Freedom of Information Act, most likely because responding is a statutory obligation. It can nonetheless still be hard to extract structured data on NHS activity from responses to FOI requests. For example, some CCGs refused to provide information on the grounds that the information requested was in the public domain (for example in web-based formularies); but we found that the public information was incomplete. CCG responses to questions can also be inconsistent, incomplete, or ambiguous. It is therefore possible that some CCG-led change activity was missed in our coding, due to CCGs not communicating it to us.

### Findings in Context

To our knowledge our key finding - that regions failing to implement change have taken little action to implement new guidance - is novel. It is also concerning. Our data is otherwise consistent with previous work describing the effectiveness of interventions to reduce inappropriate antibiotic prescribing. A Cochrane systematic review on interventions to improve antibiotic prescribing described the importance of the design of condition and situation specific interventions, including careful definition of goals; this was exhibited by four successful CCGs as per our narrative description above. The review also found that interactive educational meetings appeared to be more effective than didactic lectures; and that generic interventions, such as printed educational materials or audit and feedback alone, had little benefit.^11^ One study in Ireland found that practices with interventions including interactive workshops and decision support system prompts were more likely to prescribe nitrofurantoin first line.^12^ A Cochrane review into the use of financial incentivisation of prescribing concluded there is little evidence of benefit in the use of financial incentives to improve prescribing ^13^.

The NHS has invested extensively in “medicines optimisation” activity, where teams of pharmacists in every CCG monitor prescribing behaviour and advocate for change with individual clinicians. However there is almost no prior academic work describing the extent, character, or impact of this activity. This is surprising and concerning, given the centrality of prescribing as a medical intervention, and the importance of optimising its safety, effectiveness, and cost-effectiveness of prescribing. In 2007 The National Audit Office (NAO) asserted that value in prescribing could be improved by: communication from trusted sources and local opinion leaders; financial incentives; provision of tailored comparative (benchmarking) information to GP practices; provision of practical support such as pharmacist time to GP practices; and a coordinated approach to prescribing across the primary and secondary care sectors.^14^ These strategies have all been used, to varying degrees, in those CCGs that appeared to be most successful in promoting behaviour change among our cohort.

### Policy Implications and Interpretation

We are very concerned to see a substantial number of CCGs exhibiting minimal prescribing behaviour change, and change-related activity, in response to an important change in antibiotic prescribing policy. In our view there is room for substantial improvement in personnel training for local staff, alongside open data monitoring by NHS England, and appropriate action for those failing to implement change.

As discussed above, while there is extensive existing NHS investment in medicines optimisation staff, there is almost no literature on their activity, skills or impact, and no centralised training or curriculum. In our very extensive anecdotal experience of running an open prescribing data service with 50,000 unique users in 2017, the data skills of medicines optimisation pharmacists are highly variable. Monitoring data thoughtfully, and changing GPs’ prescribing behaviour, are challenging and specific skills. The NHS is currently investing an additional £140 million on employing 1,100 pharmacists in primary care; ^15,16^ unfortunately this has not been accompanied by a robust training programme on medicines optimisation work. In our view this is a serious oversight that should be remedied, with a training and accreditation programme for medicines optimisation.

The issue of national monitoring is also concerning. It is clear that there is some form of central monitoring around antibiotic prescribing, such as trimethoprim, if only to administer the QP payments. However we are aware of no action taken by any national NHS body when it is detected that a region or practice is prescribing anomalously; and can find no evidence in the public domain of any such action being taken. This is concerning. We suggest that such monitoring should be done openly, with all information on remedial action shared in public by default, so that others can learn from the actions undertaken.

All the structural shortcomings identified may in part be explained by the fact that responsibility for change is currently unclear. Contractual responsibility for GP quality is within the remit of NHS England; but medicines optimisation activity is currently devolved to individual CCGs. In addition, GPs retain the clinical freedom to prescribe the drugs they feel are most appropriate: while this is an important general principle, it does require that behaviour change interventions must be creative. We note that the Guillebaud Report of 1956 describes financial sanctions imposed by the NHS on individual GPs who prescribe inappropriately;^17^ to our knowledge the current GP contract would prohibit such action.

### Future Research

We suggest that alongside the development of core curricula, training, and accreditation of medicines optimisation staff there should also be basic research conducted on their current activity, skills, and impact; on the best strategies for data monitoring; and the best techniques to implement change in GP prescribing behaviour. It is surprising to find so little current evidence on the activities and impact of this large and important workforce.

### Summary

We found that many CCGs failed to implement an important change in antibiotic prescribing guidance; and report strong evidence suggesting that CCGs with minimal prescribing change did little to implement the new guidance. We strongly recommend a national programme of training and accreditation for medicines optimisation pharmacists; and remedial action for CCGs that fail to implement guidance; with all materials and data shared publicly for both such activities.

## Ethical approval

This study uses exclusively open, publicly available data; no ethical approval was required.

## Contributorship

RC and BG conceived the study. RC and BG designed the methods. RC and AW collected and analysed the data with input from BG. RC and BG drafted the manuscript. All authors contributed to and approved the final manuscript. BG supervised the project and is guarantor.

## Appendix 1: Freedom of Information Request

Dear CCG,

We are currently researching the change in GP prescribing behaviour in urinary tract infections.

Please can you provide us with the following information:

1. Do you have a formulary used by primary care prescribers? If so, what is the current status of nitrofurantoin and trimethoprim with respect to urinary tract infections (e.g. first line, second line, etc)
2. Has the status of either nitrofurantoin or trimethoprim changed in the formulary since November 2014? If so, can you please provide the previous status(es) and details of the date(s) of change.
3. Have you had any work plans in place with respect to nitrofurantoin and trimethoprim prescribing since 2014? If so, can you please provide documents and start date.
4. Have you had any GP prescribing incentive schemes or similar which relate to trimethoprim or nitrofurantoin prescribing since November 2014? If so, can you please provide the documents.

Many thanks

**Abbreviations**
BNF1: British National Formulary
CCG: Clinical Commissioning Group
CSU: Commissioning Support Unit
FOI: Freedom of Information
NAO: National Audit Office
QP: Quality Premium
UTI: Urinary-tract infection

